# The Chromatin-binding Protein Spn1 contributes to Genome Instability in *Saccharomyces cerevisiae*

**DOI:** 10.1101/341768

**Authors:** Alison K. Thurston, Catherine A. Radebaugh, Adam R. Almeida, Juan Lucas Argueso, Laurie A. Stargell

## Abstract

Cells expend a large amount of energy to maintain their DNA sequence. DNA repair pathways, cell cycle checkpoint activation, proofreading polymerases, and chromatin structure are ways in which the cell minimizes changes to the genome. During replication, the DNA damage tolerance pathway allows the replication forks to bypass damage on the template strand. This avoids prolonged replication fork stalling, which can contribute to genome instability. The DNA damage tolerance pathway includes two sub-pathways: translesion synthesis and template switch. Post-translational modification of PCNA and the histone tails, cell cycle phase, and local DNA structure have all been shown to influence sub-pathway choice. Chromatin architecture contributes to maintaining genome stability by providing physical protection of the DNA and by regulating DNA processing pathways. As such, chromatin-binding factors have been implicated in maintaining genome stability. Using *Saccharomyces cerevisiae*, we examined the role of Spn1, a chromatin binding and transcription elongation factor, in DNA damage tolerance. Expression of a mutant allele of *SPN1* results in increased resistance to the DNA damaging agent methyl methanesulfonate, lower spontaneous and damage-induced mutation rates, along with increased chronological lifespan. We attribute these effects to an increased usage of the template switch branch of the DNA damage tolerance pathway in the *spn1* strain. This provides evidence for a role of wild type Spn1 in promoting genome instability, as well as having ties to overcoming replication stress and contributing to chronological aging.

## Introduction

Maintaining the genome of a cell is of fundamental importance. Instability within the genome contributes to cancer, aging and genetic diseases (Aguilera and Garcia-Muse 2013; Vijg and Suh 2013). Point mutations, deletions, duplications, and translocations are all forms of genome instability (Aguilera and Garcia-Muse 2013; Skoneczna *et al.* 2015). Overlapping and conserved DNA repair pathways, DNA damage activated cell cycle checkpoints, proofreading DNA polymerases, and chromatin structure are some of the ways in which the cell minimizes changes to the genome (Kawasaki and Sugino 2001; Aguilera and Garcia-Muse 2013; Polo and Almouzni 2015; Chatterjee and Walker 2017). However, some level of genome instability is tolerated and is necessary to fuel genetic diversification and evolution (Skoneczna *et al.* 2015).

DNA base lesions, breaks, strand crosslinks and gaps, secondary structures and strongly bound proteins are obstacles for the replication machinery (Hustedt *et al.* 2013; Brambati *et al.* 2015; Chatterjee and Walker 2017). Replication stress caused by these structures can result in genome instability and/or cell death (Hustedt *et al.* 2013; Zeman and Cimprich 2014). Furthermore, intermediate steps of DNA damage repair can be detrimental to the cell if performed without proper coordination with replication fork progression (Ulrich 2011). For example, the cleavage of the phosphate backbone in the ssDNA template would result in a double stranded break, putting the cell at risk for chromosomal rearrangements. The DNA damage tolerance (DDT) pathway provides mechanisms for cells to circumvent blocks to the DNA replication fork (Ulrich 2011; Bi 2015; Xu *et al.* 2015; Branzei and Psakhye 2016; Branzei and Szakal 2016). DDT is different from other repair pathways since the initial damage is not immediately repaired. The DDT pathway includes two sub-pathways: the translesion synthesis branch (error prone) and a template switch branch (error free) (Lee and Myung 2008; Bi 2015; Xu *et al.* 2015; Branzei and Szakal 2016).

Translesion synthesis (TLS) utilizes polymerase switching to overcome replication blocks using the lower fidelity polymerases POL ζ (Rev3/Rev7) and Rev1 (Prakash *et al.* 2005; Xu *et al.* 2015). The TLS branch is considered error prone as it can potentially introduce a miss-matched dNTP via the low fidelity polymerase. TLS can contribute to upwards of half the point mutations accumulated by a cell at each division cycle (Stone *et al.* 2012). Template switch (the error free sub-pathway) utilizes the newly synthesized sister strand as a template for DNA synthesis past an obstruction. This requires homologous recombination factors for strand invasion and downstream DNA processing factors to resolve recombination intermediates (Branzei *et al.* 2008; Branzei and Szakal 2016). The error free sub-pathway has been determined to be different from traditional recombination repair pathways through genetic studies (Branzei and Szakal 2016). How the cell determines which DTT sub-pathway to use is still unclear; however, post-translational modification of PCNA and histone tails, cell cycle phase, and local DNA structure have all been shown to influence sub-pathway choice (Daigaku *et al.* 2010; Gonzalez-Huici *et al.* 2014; Meas *et al.* 2015; Xu *et al.* 2015; Branzei and Szakal 2016; Hung *et al.* 2017).

Spn1 (Suppresses post-recruitment gene number 1) is a transcription elongation and chromatin-binding factor (Fischbeck *et al.* 2002; Krogan *et al.* 2002; Li *et al.* 2018). *SPN1* is essential for cellular viability and is conserved from yeast to humans (Fischbeck *et al.* 2002; Liu *et al.* 2007; Pujari *et al.* 2010). Both yeast and human Spn1 consist of two intrinsically disordered tails with an ordered central core domain (Pujari *et al.* 2010) (Figure 1A) and have been shown to associate with chromatin (Kubota *et al.* 2012; Alabert *et al.* 2014; Dungrawala *et al.* 2015). Human Spn1 recruits the HYPB/Setd2 methyltransferase required for H3K36 trimethylation (H3K36me3) (Yoh *et al.* 2008). H3K36me3 through Setd2 activity has been shown to recruit homologous recombination factors to double stranded breaks (Pfister *et al.* 2014). Yeast Spn1 binds histones, DNA and nucleosomes (Li *et al.* 2018), and associates with RNA Polymerase II (RNAPII) and the histone chaperone, Spt6 (Diebold *et al.* 2010; Mcdonald *et al.* 2010; Pujari *et al.* 2010; Li *et al.* 2018). Spn1 maintains repressive chromatin in human cells (Gerard *et al.* 2015), and loss of the DNA, histone and nucleosome binding functions in yeast results in increased nucleosome occupancy at the activated *CYC1* locus (Li *et al.* 2018). Yeast Spn1 has been shown to genetically and/or physically interact with the ATP-dependent chromatin-remodelers INO80 (Costanzo *et al.* 2016) and SWR-C/SWR1 (Collins *et al.* 2007), both of which are involved in DNA double strand break repair (Van Attikum *et al.* 2007). In addition, *SPN1* genetically interacts with genes whose protein products are involved in DNA repair, such as Rad23 (Collins *et al.* 2007,Prakash and Prakash 2000), Polε (Dubarry *et al.* 2015), and histone chaperones CAF-1, Asf1 (Kim and Haber 2009; Li *et al.* 2018) and FACT (CA Radebaugh, unpublished results; Dinant *et al.* 2013). Together, these observations raise the possibility that Spn1 may influence DNA repair functions.

**Figure 1.**
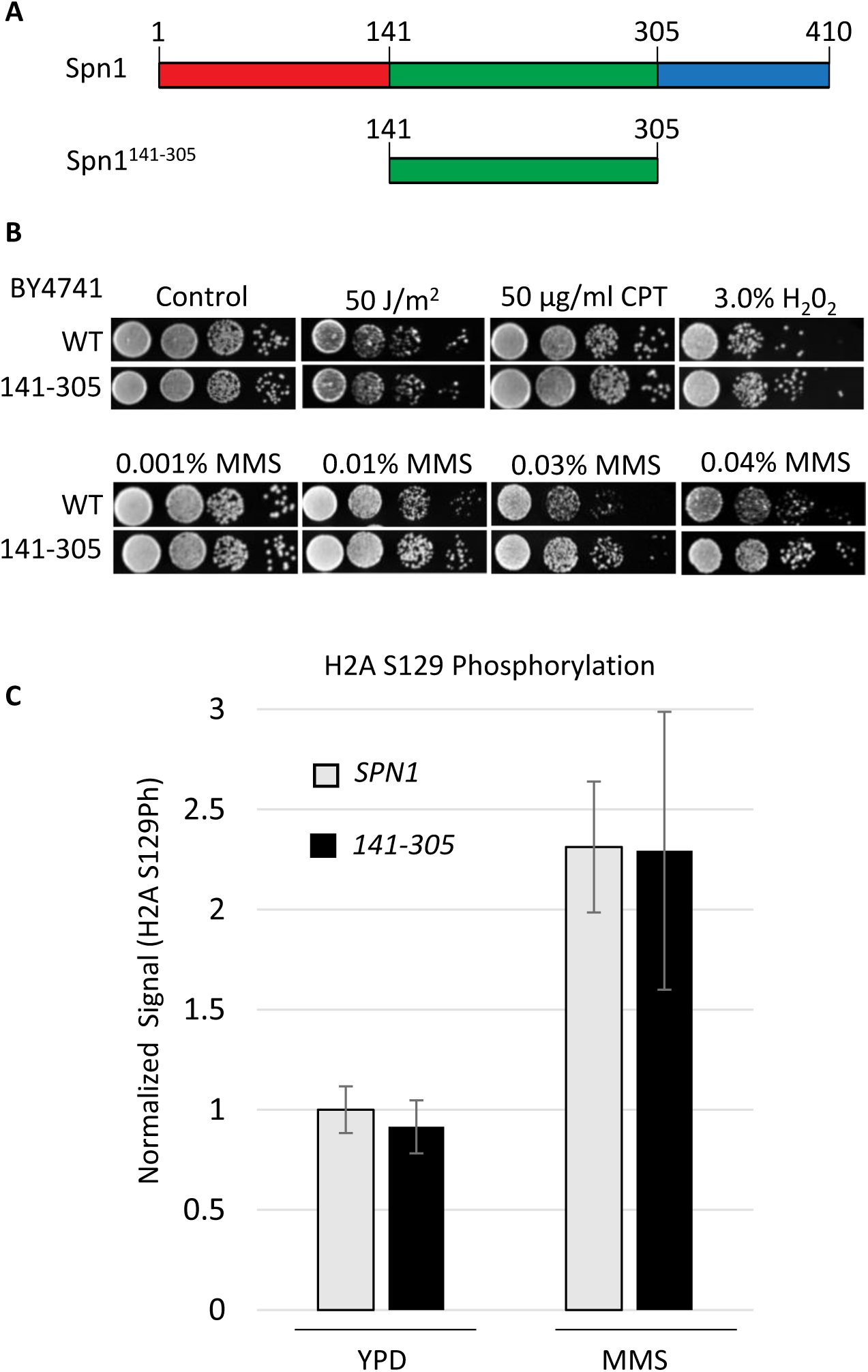
Expression of Spn1^141-305^ suppresses sensitivity to the DNA damaging agent, methyl methanesulfonate. A) Schematic of Spn1 and Spn1^141-305^. B) Ten-fold serial dilutions of cells expressing Spn1 (WT) or Spn1^141-305^ (141-305) were spotted onto YPD and YPD plates containing camptothecin (CPT), hydrogen peroxide (H2O2) or increasing concentrations of MMS. For UV damage, cells were spotted onto YPD plates and exposed to UV. C) Quantification of immunoblot showing H2A S129 phosphorylation levels before and after exposure to 0.1% MMS in cells expressing Spn1 or Spn1^141-305^. H2A S129 phosphorylation signal is normalized to TBP. All ratios are compared to wild type grown in YPD, which is set to 1. Standard deviation is calculated from 4-5 biological replicates.

In this study, we examined a role for Spn1 in genome instability in *Saccharomyces cerevisiae*. We utilized an allele, *spn1^141-305^,* which encodes a derivative that lacks the intrinsically disordered tails (Figure 1A) (Fischbeck *et al.* 2002; Li *et al.* 2018). Expression of the mutant protein Spn1^141-305^ led to increased resistance to the DNA damaging agent methyl methanesulfonate (MMS). We tested genetic interactions between *SPN1* and genes involved in base excision repair (BER), nucleotide excision repair (NER), homologous recombination (HR), and the DDT pathway. Through the analyses of these genetic interactions, the resistance to MMS observed in the *spn1^141-305^* strain was determined to be dependent on the DDT pathway and HR factors. Furthermore, the *spn1^141-305^* strain displayed decreased spontaneous and damage-induced mutation rates and increased chronological longevity. Taken together, our results indicate a role for Spn1 in promoting genome instability by influencing DNA damage tolerance sub-pathway selection.

## Materials and Methods

### Yeast culturing and strains

All strains were grown and experiments were performed in yeast peptone dextrose (YPD; 2% dextrose) media at 30°C unless otherwise indicated. A description of the yeast strains, plasmids and DNA primers utilized in this study are provided in Tables S1, S2, and S3.

The wild type strain BY4741 (*MAT***a** *his3*Δ*1 ura3*Δ*0 leu2*Δ*0 met15*Δ*0*) and deletion strains were purchased from Thermo Scientific Open. Strains were created as described (Zhang *et al.* 2008; Li *et al.* 2018). Briefly, strains were transformed with a covering plasmid expressing Spn1. Endogenous *SPN1* was replaced by a *LEU2* fragment flanked by *SPN1* promoter sequences by homologous recombination. Plasmids containing *SPN1* or *spn1^141-305^* were introduced into strains by plasmid shuffling.

### DNA damage exposure phenotypic assays

To assess the growth phenotypes and possible genetic interactions between *SPN1* and other mutant strains, yeast cells were grown overnight in YPD. Cultures were diluted, grown to log phase, collected by centrifugation, washed with sterile water and diluted. Ten-fold serial dilutions were spotted onto the indicated solid media. Plates were grown at 30°C. Images of plates were taken daily. Methyl methanesulfonate (MMS), camptothecin (CPT) and hydrogen peroxide (H_2_O_2_) plates were made 24-48 hours before each experiment. UV exposure was performed with a UVP UVLMS-38 light source at a wavelength of 254 nm. To test if *SPN1* is dominant, strains were grown using SC-His drop-out media to maintain selection of the plasmid expressing Spn1 or Spn1^141-305^.

### Fluctuation analysis

Fluctuation analyses were performed to determine the rates of spontaneous and damage-induced forward mutation in the *CAN1* gene. Indicated strains were patched and grown for 24 hours on YPD. Strains were streaked to single colonies onto YPD plates and grown for 48 hours. Multiple colonies of each strain were inoculated and allowed to grow for 24 hours in 5 mL of YPD. Cells were pelleted and washed in sterile water, and appropriate dilutions were plated on YPD and SC-Arg drop-out containing 60 μg/L of canavanine. Colonies were counted after two (YPD, permissive) and three (SC-Arg drop-out + Can, selective) day growth. Mutation rates were calculated through the Lea-Coulson method of the median (Lea and Coulson 1949) in using the FALCOR web application (Hall *et al.* 2009). Statistical significance was determined using the Mann-Whitney non-parametric t-test using GraphPad Prism software. For damage-induced mutation rates, the same protocol was followed except strains were streaked onto YPD plates containing 0.001% MMS and inoculated into liquid YPD containing 0.005% MMS. *mms2Δ* mutants were inoculated in YPD liquid containing 0.001% MMS due to this strain’s higher sensitivity to MMS. Plates containing MMS were made less than 24 hours before use.

### Budding index

Strains were grown overnight in YPD media, cultures were then diluted and grown to log phase. The cultures were split and MMS was added to half of the cells for a final concentration of 0.03% MMS. Cultures were incubated for an additional 30 minutes at 30°C. The cells were washed and fixed with formalin. At least 300 cells were evaluated for each strain; determination of cell cycle phase was determined by bud size analysis.

### Immunoblotting analysis

Cells were harvested at log phase and suspended in 0.1 M NaOH for 5 minutes. The NaOH was removed, the cell pellet was resuspended in lysis buffer (120 mM Tris-HCl [pH 6.8], 12% glycerol, 3.4% SDS, 200 mM dithiothreitol [DTT], 0.004% bromophenol blue), and the cell suspension was incubated at 95°C for 5 minutes. Insoluble cell debris was removed by centrifugation, and total protein was separated on a 15% SDS-PAGE gel. Proteins were transferred to a nitrocellulose blotting membrane (Amershan Protran 0.2uM NC) and blocked with 3% milk. The following antibodies were utilized: anti-TBP (1:5,000), anti-H2AS129 phosphorylation (abcam #ab15083, 1:500), and anti-rabbit (Li-COR #925-32211, 1:15000). Protein bands were visualized using the Li-COR Odyssey CLx and quantified using Image Studio.

### Chronological aging assay

Chronological aging assays were performed as described (Parrella and Longo 2008). Briefly, strains were inoculated in synthetic dropout media and grown overnight. Cultures were diluted to an OD of 0.1 and grown in synthetic dropout media for 3 days (72 hours) to ensure cultures have reached stationary phase (**T_0_**). To determine viability, dilutions of each biological replicate were plated every other day onto YPD plates. Dilutions and plating were carried out in triplicate and averaged for each biological replicate. Four to five biological replicates for each strain were averaged to determine the % viability. The % viability is the ratio of viable colonies at a specific time (**T_x_**) over the number of viable colonies at **T_0_** (stationary phase).

### Data Availability

Strains and plasmids are available upon request. The authors affirm that all data necessary for confirming the conclusions of the article are present within the article, figures, tables and supplemental material. Supplemental data is available at Figshare as one PDF. Figure S1 shows further analysis of the MMS resistance. Figure S2 examines growth phenotypes after UV exposure. Figure S3 examines growth phenotypes after exposure to MMS or HU in DDT deletion background strains. Figure S4 compares DDT sub-pathway selection between wild type and the spn1141-305 strain. Table S1, Table S2 and Table S3 list the strains, plasmids and DNA primers utilized in this study, respectively.

## Results

### Expression of Spn1^141-305^ results in increased resistance to methyl methanesulfonate

To test if Spn1 functions in DNA damage repair, we utilized the *spn1* mutant allele *spn1^141-305^* (Figure 1A). The Spn1^141-305^ protein consists of only the conserved ordered core domain, and lacks the N and C termini present in full length Spn1 (Li *et al.* 2018). Cells expressing Spn1^141-305^ were grown on media containing various DNA damaging agents and interestingly, the *spn1^141-305^* strain displayed resistance to MMS (Figure 1B). The observed resistance is specific to MMS since sensitivity to the other DNA damaging agents was not altered (Figure 1B). Co-expression of plasmid-borne Spn1^141-305^ in a strain carrying a chromosomal copy of wild type Spn1 did not result in increased resistance to MMS (Figure S1A), suggesting that *spn1^141-305^* is recessive.

To determine whether cells expressing Spn1^141-305^ are able to trigger a DNA damage response after exposure to MMS, the levels of histone H2A serine 129 phosphorylation (H2A S129Ph) in *SPN1* and *spn1^141-305^* cells were measured by immunoblot analysis. H2A S129 is phosphorylated in response to DNA damage (Downs *et al.* 2000; Foster and Downs 2005). A ~2.5 fold increase in the amount of H2A S129Ph after MMS exposure was observed in both the *SPN1* and *spn1^141-305^* strains (Figure 1C). The levels of H2A S129Ph were similar in the two strains prior to and after exposure to MMS (Figure 1C). In addition, we examined the cell cycle phase distribution between the two strains after exposure to MMS (Figure S1B). No difference was observed between the cell cycle phase distributions of the two strains. Since, both the H2A S129Ph DNA damage response and cell cycle progression were not altered in the *spn1^141-305^* strain, we reasoned that the DNA damage checkpoints remained intact.

Mec1 and Tel1 are evolutionarily conserved phosphatidylinositol-3 kinase related protein kinases (PIKKs) and transduce a kinase cascade, which activates DNA damage repair, cell cycle arrest, transcription programs, dNTP synthesis, and replication fork stabilization in response to cellular stress (Craven *et al.* 2002; Toh and Lowndes 2003; Enserink 2011). Serine 23 (S23) of Spn1 is phosphorylated in response to exposure to MMS in a Mec1- and Tel1-dependent and Rad53-independent manner (Chen *et al.* 2010; Bastos de Oliveira *et al.* 2015). We investigated whether cells expressing Spn1^141-305^ combined with the loss of Tel1 or Rad9, a mediator kinase that activates Rad53, remain MMS resistant. We observed loss of resistance when either Tel1 or Rad9 were deleted (Figure S1C). Since Tel1 and Rad9 affect many downstream factors we wanted to test if the loss of Spn1 S23 phosphorylation is sufficient for MMS resistance. Phospho-mimetic (*spn1^S23D^*) and phospho-deficient (*spn1^S23A^*) strains were created and grown on MMS. The *spn1^S23D^* and *spn1^S23A^* strains grew similarly to the *SPN1* strain (Figure S1D). Taken together, this indicates that MMS resistance observed in the *spn1^141-305^* strain is dependent on Tel1 and Rad9 activity and that the loss of S23 phosphorylation of Spn1 is insufficient for MMS resistance.

### Removal of methyl lesions through Mag1 glycosylase is necessary for MMS resistance

To investigate if the resistance to MMS could be due to more efficient DNA repair, we introduced *spn1^141-305^* into strains deficient for base excision repair. BER is the primary pathway to repair damage caused by MMS (Memisoglu and Samson 2000). Mag1 is the DNA glycosylase responsible for the removal of the toxic N3-methyladenine adducts resulting in an abasic site (Prakash and Prakash 1977; Chen *et al.* 1989). Apn1 is the major endonuclease responsible for cleavage of the phosphate backbone at the abasic site, which is subsequently repaired through long or short patch BER (Memisoglu and Samson 2000; Odell *et al.* 2013). Cells expressing either Spn1 or Spn1^141-305^ in the *mag1Δ* background were sensitive to MMS (Figure 2A). In contrast, cells expressing Spn1^141-305^ in the *apn1Δ* background were resistant to MMS (Figure 2A). This indicates that cells bearing *spn1^141-305^* are able to retain resistance with a defective BER pathway, if the damaging lesion can be processed by Mag1.

**Figure 2.**
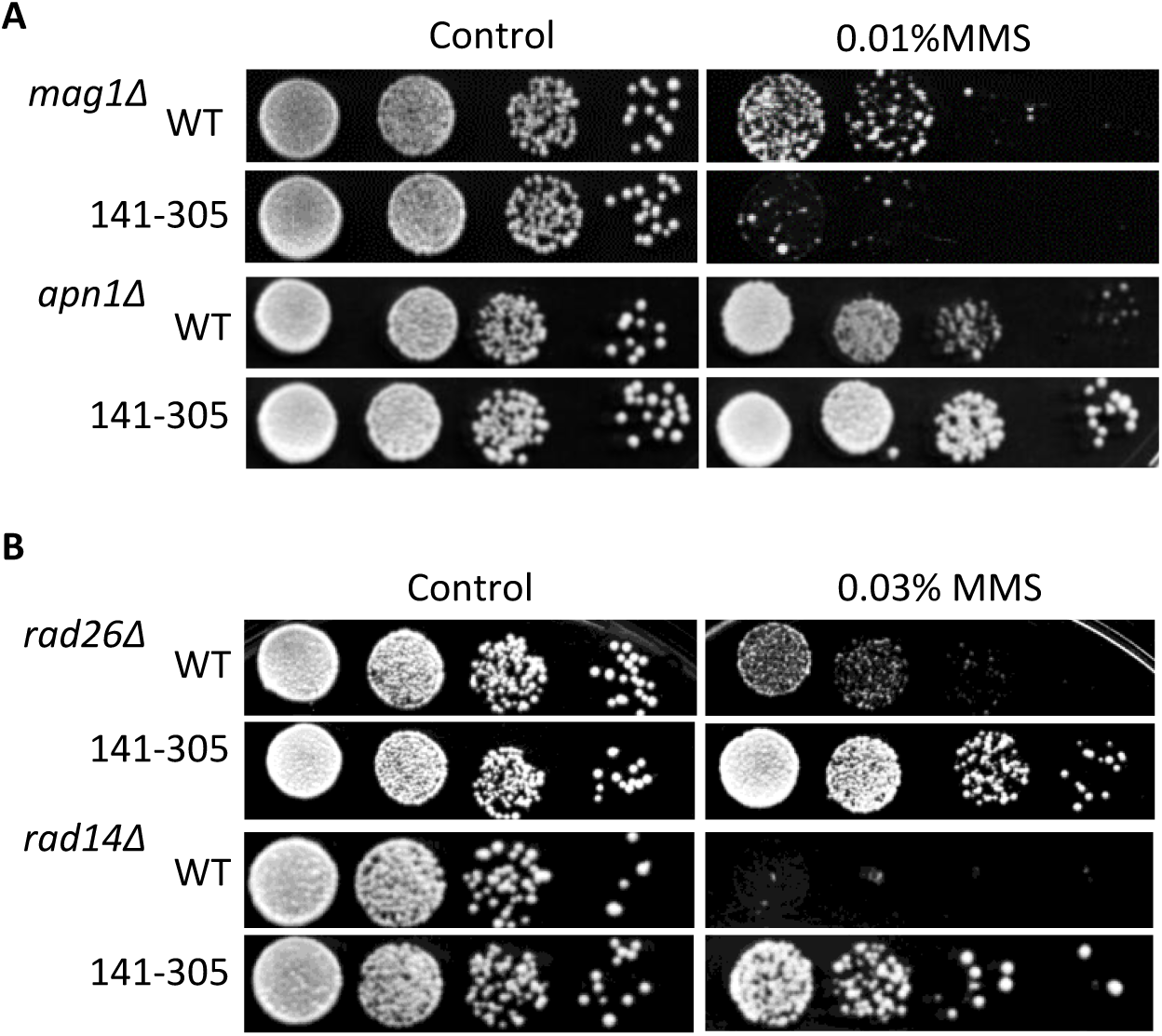
Resistance to MMS is dependent on a functional BER pathway and independent of NER. Ten-fold serial dilutions of cells expressing Spn1 (WT) or Spn1^141-305^ (141-305) in A) *mag1Δ* nd *apn1Δ* backgrounds and B) *rad26Δ* and *rad14Δ* backgrounds. Cells were grown on YPD and MMS plates.

### Resistance to MMS is independent of the nucleotide excision repair pathway

Since Spn1 is involved in transcription and mRNA processing (Fischbeck *et al.* 2002; Krogan *et al.* 2002; Yoh *et al.* 2007; Yoh *et al.* 2008), it seemed possible that Spn1 could be functioning in transcription-coupled nucleotide excision repair (TC-NER). Additionally, nucleotide excision repair (NER) has been show to compete with BER AP endonucleases in the repair of abasic sites (Torres-Ramos *et al.* 2000). Genetic analyses were performed with the introduction of *spn1^141-305^* into the *rad26Δ* and *rad14Δ* backgrounds. Rad26 is a DNA-dependent ATPase involved in TC-NER (Guzder *et al.* 1996a; Prakash and Prakash 2000), and Rad14 is a subunit of NER factor 1 (NEF1), which is required for TC-NER and global genomic NER (Guzder *et al.* 1996b; Prakash and Prakash 2000). Resistance to MMS was maintained when Spn1^141-305^ was expressed in the *rad26Δ* or the *rad14Δ* strain (Figure 2B), indicating that this phenotype is not dependent on either NER pathway. Furthermore, expression of Spn1^141-305^ in the wild type, *rad26Δ* and *rad14Δ* strain backgrounds did not result in a resistance phenotype after exposure to UV (Figure S2A and S2B), which is primarily repaired by NER. This further supports that the resistance phenotype is specific to MMS and it is not due to enhancement of the NER pathway.

### Resistance is dependent on the error free sub-pathway of the DNA damage tolerance pathway

The DDT pathway provides a mechanism for cells to bypass blocks to the DNA replication fork (Ulrich 2011; Bi 2015), including lesions caused by exposure to MMS (Figure S3A). The primary signal for entry into the TLS sub-pathway is dependent on the mono-ubiquitination of PCNA through the actions of the Rad18/Rad6 complex (Ulrich 2011; Bi 2015). Additional polyubiquitination through the actions of the Ubc13/Mms2 and Rad5 complex is the primary signal for error free bypass (Ulrich 2011; Bi 2015). It has been shown that abolishment of the DDT pathway results in extreme sensitivity to MMS (Huang *et al.* 2013). Since we observe resistance to MMS in the *spn1^141-305^* strain, we predicted that the DDT pathway must function. Consistent with this, deletion of *RAD6* or *RAD18* resulted in the loss of MMS resistance in strains expressing Spn1^141-305^ (Figure S3B).

MMS resistance has been shown to correlate with changes in DDT sub-pathway selection (Conde and San-Segundo 2008; Conde *et al.* 2010). Genetic analyses between *SPN1* and *REV3/REV7/REV1*, TLS polymerases, and *RAD5/MMS2/UBC13*, a complex responsible for error free sub-pathway signaling, were performed. Cells expressing Spn1^141-305^ retained resistance to MMS in all tested TLS gene deletion backgrounds (Figure 3A). In contrast, loss of MMS resistance was observed in the error-free deletion strains (Figure 3B). These data suggest cells expressing wild type Spn1 are utilizing the TLS sub-pathway, whereas cells expressing Spn1^141-305^ shift the response toward the error free sub-pathway.

**Figure 3.**
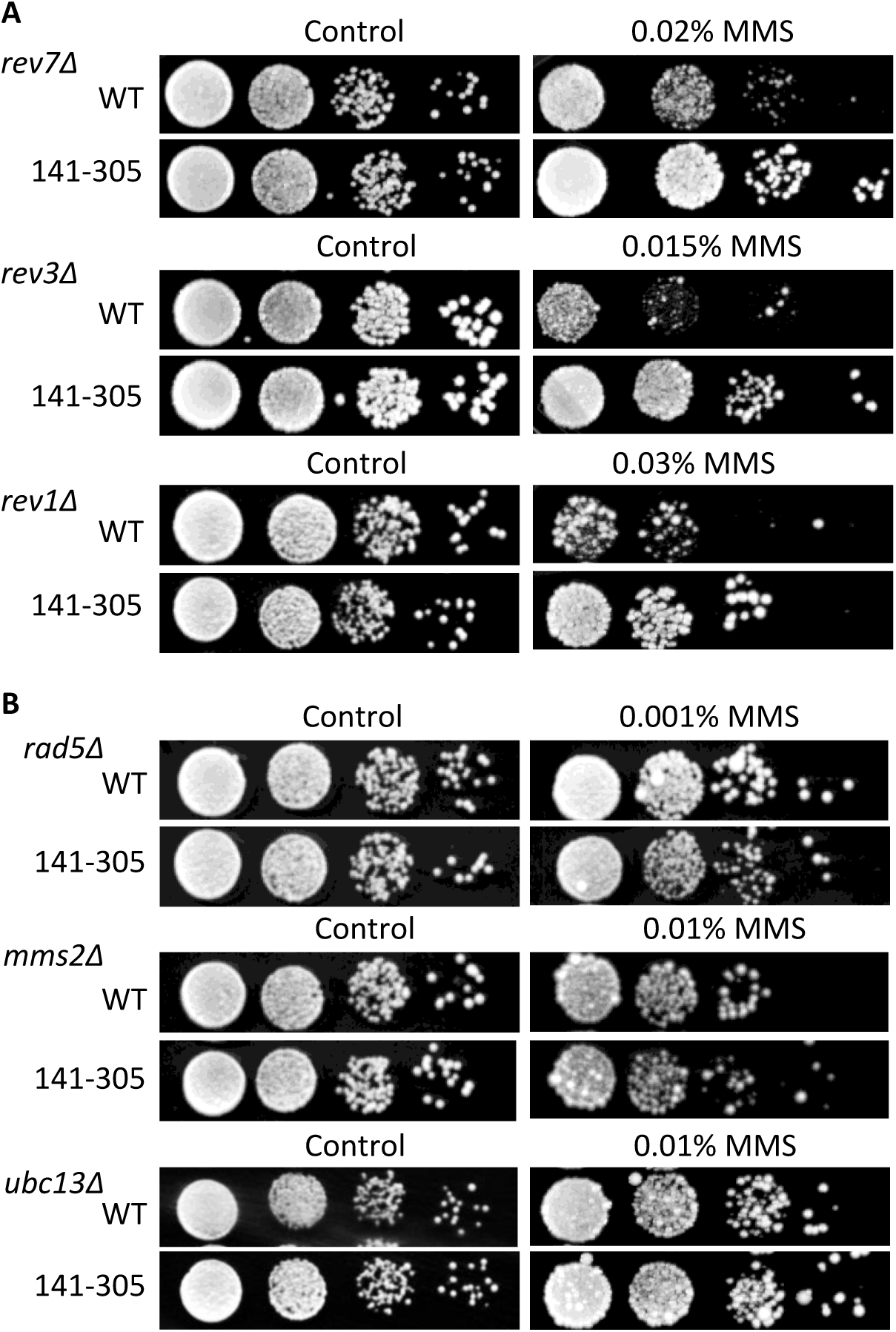
Resistance to MMS is dependent on the error free branch of the DNA damage tolerance pathway. Ten-fold serial dilutions of cells expressing Spn1 (WT) or Spn1^141-305^ (141-305) in A) TLS deletion background strains and B) error free deletion background strains. Strains were grown and spotted onto YPD and MMS plates.

To test if expression of Spn1^141-305^ confers resistance to other forms of replication stress, we examined cellular growth after the addition of hydroxyurea (HU). Exposure to HU results in slowed or stalled replication forks due to depleted levels of dNTPs (Koc *et al.* 2004), while MMS causes damage-induced replication stress (Wyatt and Pittman 2006). In contrast to resistance to MMS, the *spn1^141-305^* strain displayed a slight sensitivity to HU (Figure S3C). HU sensitivity was exacerbated in the DDT deletion strains (Figure S3C). This suggests that Spn1 is important for overcoming replication stress caused by HU and that the stress induced by HU cannot be overcome by expression of Spn1^141-305^.

### Spn1 contributes to spontaneous and damage-induced genome instability

The TLS polymerases can be responsible for upwards of 50% of point mutations in a genome (Stone *et al.* 2012). Thus, we predicted that if the *spn1^141-305^* strain does not utilize the TLS sub-pathway, then we would observe a difference in the damage-induced mutation rates between the *SPN1* and *spn1^141-305^* strains. To determine levels of damage-induced mutations, a fluctuation assay detecting forward mutations occurring within the *CAN1* locus in the presence of MMS was performed. Cells expressing Spn1^141-305^ had a significant decrease in the damage-induced mutation rate compared to wild type cells (Table 1). The *spn1^141-305^* strain also displayed a decrease in the spontaneous mutation rate (Table 1). These results indicate that Spn1 contributes to genome instability, regardless of the presence of a DNA damaging agent.

**TABLE 1.** Damage induced and spontaneous mutation rates of cells expressing Spn1 (WT) or Spn1^141-305^ (141-305) in *rev3Δ, mms2Δ* and *dot1Δ* strains.

| MMS Induced Mutation Rate |  |  |  |  |  |
| --- | --- | --- | --- | --- | --- |
| Strain | Mutation Rate<sup>a</sup><br>x 10<sup>-7</sup> | 95% CI | Fold Change<sup>b</sup> | Number of Replicates | One Tailed<sup>c</sup> |
| *SPN1* | 22.76 | 19.12 - 25.83 | 1X | 21 |  |
| *spn1*<sup>141-305</sup> | 15.46 | 12.4 - 15.98 | 0.68 | 21 | <0.0001 |
| *rev3Δ* | 8.42 | 15.09 - 4.61 | 1X | 20 |  |
| *rev3Δ spn1*<sup>141-305</sup> | 5.06 | 3.66 - 5.89 | 0.60 | 21 | 0.0041 |
| *mms2Δ* | 289.63 | 250.24 - 391.05 | 1X | 21 |  |
| *mms2Δ spn1*<sup>141-305</sup> | 240.62 | 180.9 - 334.16 | 0.83 | 21 | 0.0815 |
| *dot1Δ* | 90.01 | 77.06 - 102.78 | 1X | 14 |  |
| *dot1Δ spn1*<sup>141-305</sup> | 63.13 | 58.29 - 67.8 | 0.70 | 13 | 0.0002 |
| Spontaneous Mutation Rate |  |  |  |  |  |
| Strain | Mutation Rate<sup>a</sup><br>x 10<sup>-7</sup> | 95% CI | Fold Change<sup>b</sup> | Number of Replicates | One Tailed<sup>c</sup> |
| *SPN1* | 1.2 | 0.96 - 1.59 | 1X | 21 |  |
| *spn1*<sup>141-305</sup> | 0.64 | 0.44 - 0.84 | 0.53 | 21 | <0.0001 |
| *rev3Δ* | 2.02 | 1.36 - 2.49 | 1X | 7 |  |
| *rev3Δ spn1*<sup>141-305</sup> | 0.95 | 0.65 - 3.57 | 0.47 | 7 | 0.0189 |
| *mms2Δ* | 24.29 | 13.98 - 27.97 | 1X | 14 |  |
| *mms2Δ spn1*<sup>141-305</sup> | 26 | 12.12 - 37.17 | 1.07 | 14 | 0.1344 |
| *dot1Δ* | 2.86 | 1.53 - 7.23 | 1X | 7 |  |
| *dot1Δ spn1*<sup>141-305</sup> | 2.06 | 1.73 - 2.63 | 0.72 | 7 | 0.0318 |

<sup>a</sup> Mutation rates were calculated through the Lea-Coulson method of the median (LEA and COULSON 1949) in using the FALCOR web application (HALL *et al.* 2009).
<sup>b</sup> Reported fold change is the comparison of the mutation rate of cells expressing Spn1<sup>141-305</sup> compared to wild type Spn1 in the corresponding strain background.
<sup>c</sup> Statistical significance was determined using the Mann-Whitney non-parametric t-test using GraphPad Prism software.

To test if the decreased mutation rate observed in the *spn1^141-305^* strain is dependent on the error free sub-pathway, damage-induced mutation rates in *mms2Δ* were examined. We predicted that introduction of Spn1^141-305^ into the *mms2Δ* strain would result in the mutation rate increasing and returning to levels observed in *mms2Δ* expressing wild type Spn1. As predicted, the deletion of *MMS2* combined with the *SPN1* mutant resulted in wild type damaged induced mutation rates (Table 1). Furthermore, deletion of *rev3Δ* (a component of the TLS pathway) in the *spn1^141-305^* strain still produced a decrease in mutation rate (Table 1). Taken together, this indicates that the mutation rate decrease in the *spn1^141-305^* strain is dependent on the error free sub-pathway.

The deletion of the histone methyltransferase Dot1 results in resistance to MMS through the loss of inhibition of the TLS sub-pathway (Conde and San-Segundo 2008). The resistance in the *spn1^141-305^* strain is due to use of the error free sub-pathway (Figure S3A), and thus we predicted that Spn1 and Dot1 are acting in parallel pathways. To test this, a genetic analysis between *SPN1* and *DOT1* was performed. Interestingly, combining the deletion of *DOT1* with *spn1^141-305^* resulted in increased growth compared to *dot1Δ* alone on YPD (Figure S3D). This increased growth was enhanced when cells were grown on plates containing MMS (Figure S3D). Expression of Spn1*^141-305^* in the *dot1Δ* strain resulted in significantly decreased mutation rates (Table 1), although we observed higher damage-induced mutation rates when *DOT1* was deleted, which is consistent with previously reported data (Conde and San-Segundo 2008). The increase in MMS resistance and the mutation rate observed in the *dot1Δ spn1^141-305^* strain suggests a deregulation of both sub-pathways of DDT.

### Resistance to MMS is dependent on homologous recombination machinery

The template switching mechanism utilized in the error free sub-pathway requires many of the factors involved in homologous recombination (HR) (Branzei *et al.* 2008; Branzei and Szakal 2016; Hanamshet *et al.* 2016), thus we predicted the MMS resistance seen in the *spn1^141-305^* strain would require various HR factors. During DDT, ssDNA resulting from re-priming of the replication fork is bound by Rad51 (Gangavarapu *et al.* 2007; Symington *et al.* 2014). Rad55 and Rad57 work as a heterodimer to stabilize the association of Rad51 with the ssDNA (Symington *et al.* 2014). Deletion of *RAD51*, *RAD55* or *RAD57* combined with *spn1^141-305^* resulted in loss of MMS resistance (Figure 4A), indicating the observed resistance to MMS is dependent on functional HR factors.

**Figure 4.**
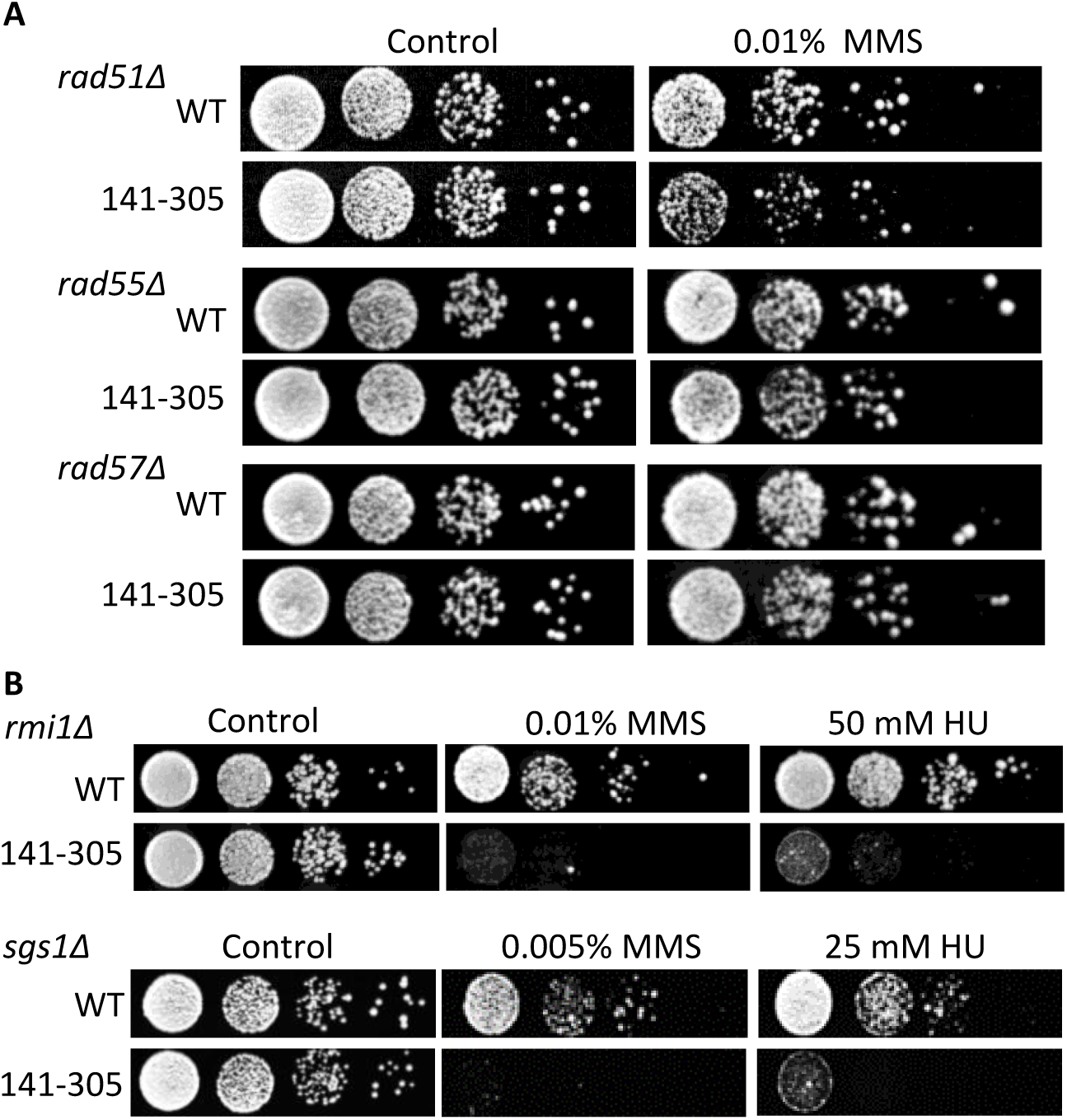
Resistance to MMS is dependent on the homologous recombination factors. Tenfold serial dilutions of cells expressing Spn1 (WT) or Spn1^141-305^ (141-305) in A) *rad51Δ*, *rad55Δ* and *rad57Δ* backgrounds and B) *sgs1Δ* and *rmi1Δ* backgrounds. Strains spotted on YPD and YPD plates containing MMS or HU.

### DNA intermediates are processed through Sgs1 and Rmi1 in spn1^141-305^

During error free DDT and HR, DNA crossover intermediates are a result of strand invasion. The functions of topoisomerases, helicases, and nucleases aid in resolving these intermediates (Mitchel *et al.* 2013; Campos-Doerfler *et al.* 2018). Sgs1, Rmi1 and Top3 work as a complex to aid in resolving holiday junctions (Mullen *et al.* 2005; Bernstein *et al.* 2009). Genetic analysis revealed that the deletion of *SGS1* or *RMI1* is synthetically lethal with *spn1^141-305^* on MMS (Figure 4B). This was also observed when the strains were grown on HU (Figure 4B). Together these observations suggest that cells expressing Spn1^141-305^ may commit to recombination pathways, and are therefore dependent on a functional Sgs1/Rmi1 complex to resolve recombination intermediates.

### Spn1^141-305^ expression results in increased chronological longevity

Decreased mutation rates and inactivation of the TLS pathway have been linked to increased chronological longevity (Longo and Fabrizio 2012). Since cells expressing Spn1^141-305^ have decreased mutation rates, we asked whether they would have an increased chronological lifespan. A dramatic difference in the chronological lifespan between cells expressing Spn1 and Spn1^141-305^ was observed (Figure 5). At the termination of the assay (19 days), the *spn1^141-305^* culture maintained 85% viability, while the control *SPN1* culture was around 5%. This suggests a link between Spn1, genome instability and chronological aging.

**Figure 5:**
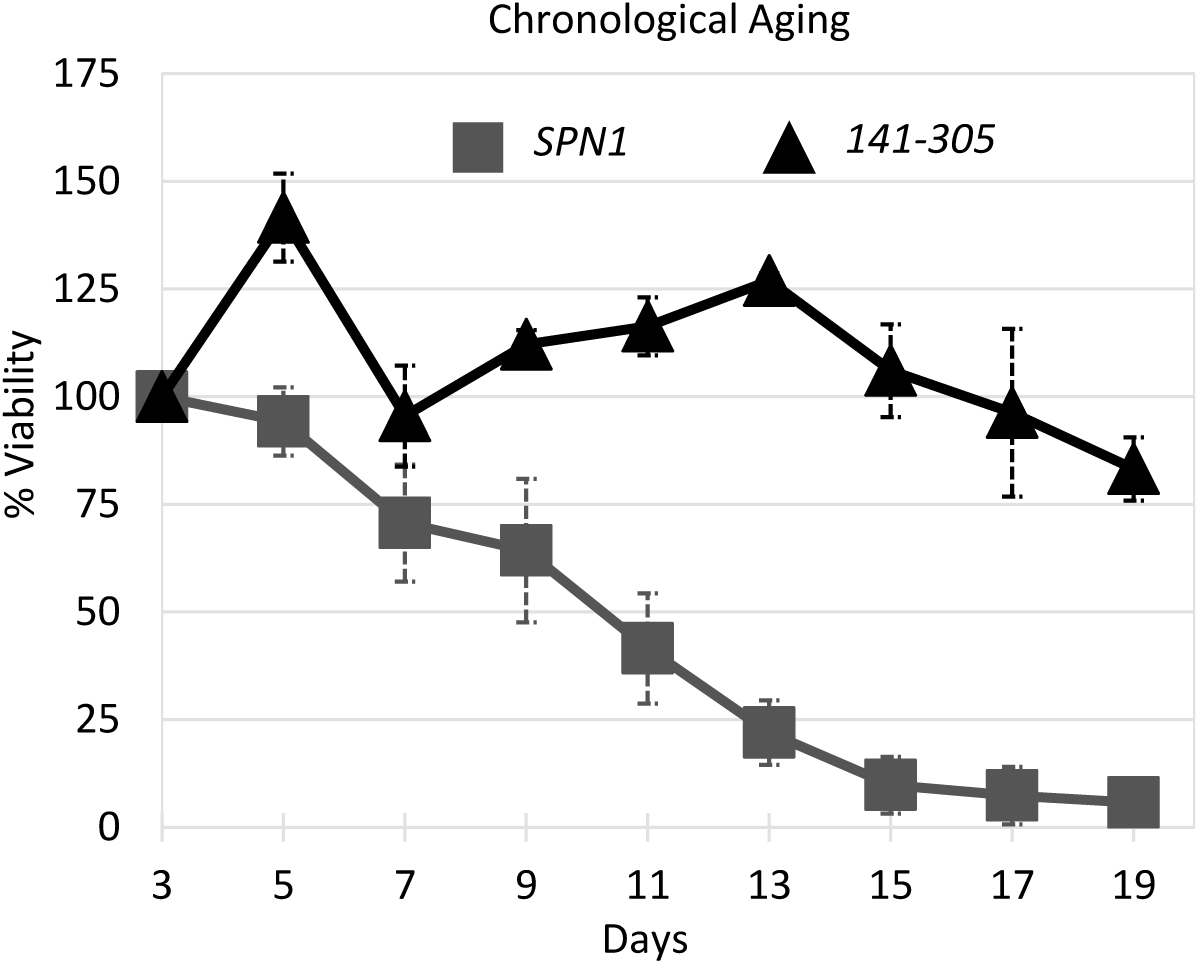
Expression of spn1^141-305^ increases chronological life span. Each time point represents the average cell viability at the number of days indicated for *SPN1* (squares; n=5) and *spn1^141-305^* (triangles; n=4).

## Discussion

Here we have investigated the role of the chromatin-binding factor Spn1 in DNA damage response and genome instability. Expression of Spn1^141-305^ covers the essential functions of wild type when cells are grown in rich culturing conditions, although this derivative has lost the ability to bind DNA, nucleosomes and histones (Li *et al.* 2018). Upon exposure to the DNA damaging agent MMS, we observed increased resistance in cells expressing Spn1^141-305^. MMS results in methylation of single and double stranded DNA (Yang *et al.* 2010). The methyl group is primarily transferred to a double bonded nitrogen on adenine, cytosine and guanine with varying frequencies (Wyatt and Pittman 2006). Not all lesions induced by MMS are toxic; however, N3-methyladenine is toxic to cells as it creates a barrier for the replication machinery (Chang *et al.* 2002). A DNA damage response was detected after exposure to MMS in both the wild type (*SPN1*) and the *spn1^141-305^* strain. DNA lesions caused by MMS are primarily repaired through BER, although other repair pathways such as NER can partially compensate (Bauer *et al.* 2015). Deletion of *MAG1*, the DNA glycosylase responsible for the recognition and removal of the toxic N3-methyladenine, results in cell sensitivity to MMS (Prakash and Prakash 1977). Expression of Spn1^141-305^ in the *mag1Δ* strain could not suppress the MMS sensitivity observed in the *mag1Δ* strain, implying Mag1 activity is necessary for resistance to MMS. Interestingly, cells expressing Spn1^141-305^ retain resistance in the *apn1Δ* strain. We reasoned that the initial removal of the methylated base is necessary for MMS resistance. Once Mag1 removes the affected base, the resulting abasic site could be processed by other endonucleases in BER or overlapping DNA repair pathways. One such pathway is NER, and it seemed plausible that Spn1 might function in NER, especially transcription-coupled: Spn1 has been shown to function as a transcription elongation factor and has physical and genetic interactions with transcription factors and RNA Polymerase II (Fischbeck *et al.* 2002; Krogan *et al.* 2002; Pujari *et al.* 2010). Loss of Rad14 (a NER factor) is lethal when cells are exposed to MMS or UV. Although expression of Spn1^141-305^ in the *rad14Δ* strain suppressed cell death when grown on MMS, the expression of Spn1^141-305^ (*rad14Δ spn1^141-305^*) could not rescue lethality after exposure to UV. Additionally, the *spn1^141-305^* strain revealed no mutant UV phenotype, indicating that expression of Spn1^141-305^ was not enhancing NER repair.

Further genetic analysis revealed that the resistance observed in the *spn1^141-305^* strain was dependent on the error free sub-pathway of DDT. Resistance was lost upon deletion of any of the genes responsible for poly-ubiquitination of PCNA (*RAD5/MMS2/UBC13*), the major signal for entry into the error free sub-pathway. Error free bypass utilizes HR factors for template switching (Branzei and Szakal 2016; Hanamshet *et al.* 2016). We observe loss of resistance in all genes tested in the HR group (*RAD51/RAD55/RAD57*). Template switching requires additional downstream HR factors to resolve DNA intermediates. Introduction of *spn1^141-305^* into the *sgs1Δ* or *rmi1Δ* strain is synthetically lethal when grown on MMS and HU. We conclude that expression of Spn1^141-305^ shifts the regulation of DDT towards the error free sub-pathway (Figure S4) and resolution of the resulting DNA intermediates is dependent on the function of the Sgs1/Rmi1/Top3 complex. This shift in regulation of the DDT pathway appears advantageous after exposure to MMS, however when cells are exposed to HU, the effects of expression of Spn1^141-305^ are detrimental to cell growth. This could be due to the amount of replication stress a cell is experiencing during exposure to MMS versus HU. These results indicate a role for wild type Spn1 in overcoming replication stress caused by HU and utilizing the TLS sub-pathway.

In cells expressing Spn1^141-305^, the shift towards the error-free pathway results in a significant decrease in genome instability. The decrease is lost upon deletion of *MMS2* in cells expressing Spn1^141-305^, indicating the decreased mutation rates observed are dependent on the error free sub-pathway of DDT. This is intriguing because it indicates wild type Spn1 contributes to genome instability.

In addition to decreased mutation rates in the *spn1^141-305^* strain, we also observed increased chronological longevity. As yeast age, the frequency of all types of mutations increases (Madia *et al.* 2007; Longo and Fabrizio 2012). Decreases in accumulated mutations are linked to a cell’s ability to process damaged DNA (primarily oxidative damage), decrease activity of the TLS polymerases, control mitotic recombination rates and regulate metabolism (Madia *et al.* 2009). Cells lacking Sch9, a protein kinase, resulted in increased chronological longevity (Longo and Fabrizio 2012), which is linked to the inactivation of the Rev1-Polζ polymerases and decreased damage accumulation (Madia *et al.* 2009; Longo and Fabrizio 2012). Decreased dependence on the TLS sub-pathway resulting in decreased mutation accumulation could contribute to the increased chronological lifespan observed in cells expressing Spn1^141-305^.

Our results show that Spn1 influences the DDT pathway, however a specific mechanism remains to be determined. Chromatin structure, histone tail modification, DNA topography, and DNA sequence all influence DDT pathway selection (Gonzalez-Huici *et al.* 2014; Meas *et al.* 2015; Hung *et al.* 2017). Deletion of the H3K79 methyltransferase Dot1 also results in resistance to MMS (Conde and San-Segundo 2008). The loss of TLS inhibition resulting in MMS resistance in *dot1Δ* was determined to be due to the loss of the methylase activity (Conde *et al.* 2010). Interestingly, increased MMS resistance is observed in the *dot1Δ spn1^141-305^* strain suggesting deregulation of both sub-pathways of DDT, possibly due to aberrant chromatin structure. Chromatin in cells expressing Spn1^141-305^ is more resistant to micrococcal nuclease digestion during activated *CYC1* transcription than wild type cells (Li *et al.* 2018), and Spn1 prevents the chromatin remodeler SWI/SNF from being recruited during repression of *CYC1* transcription (Zhang *et al.* 2008). Both human and yeast Spn1 associate with chromatin throughout the cell cycle (Kubota *et al.* 2012; Alabert *et al.* 2014; Dungrawala *et al.* 2015), and human Spn1 was determined to be an early arriving chromatin component factor after DNA was replicated (Alabert *et al.* 2014). Thus, one possibility is that wild type Spn1, which is capable of binding DNA, histone, and nucleosomes, aids in the creation and/or maintenance of a chromatin architecture that tips the balance between error-prone and error-free DDT sub-pathway utilization.

## Acknowledgements

We thank Tyler Glover, Sarah Stonedahl, Rachael Carstens and Raira Ank for general lab assistance and cell culturing. We thank Nadia Sampaio and Alexandra Gehring for technical assistance, and Thomas Santangelo and Eric Ross for reagents and equipment. This work was supported by National Science Foundation [MCB-1330019 to L.A.S.].

## Literature Cited

Aguilera, A., and T. Garcia-Muse, 2013 Causes of genome instability. Annu Rev Genet 47: 1–32.

Alabert, C., J. C. Bukowski-Wills, S. B. Lee, G. Kustatscher, K. Nakamura et al., 2014 Nascent chromatin capture proteomics determines chromatin dynamics during DNA replication and identifies unknown fork components. Nat Cell Biol 16: 281–293.

Bauer, N. C., A. H. Corbett and P. W. Doetsch, 2015 The current state of eukaryotic DNA base damage and repair. Nucleic Acids Res 43: 10083–10101.

Bernstein, K. A., E. Shor, I. Sunjevaric, M. Fumasoni, R. C. Burgess et al., 2009 Sgs1 function in the repair of DNA replication intermediates is separable from its role in homologous recombinational repair. EMBO J 28: 915–925.

Bi, X., 2015 Mechanism of DNA damage tolerance. World J Biol Chem 6: 48–56.

Brambati, A., A. Colosio, L. Zardoni, L. Galanti and G. Liberi, 2015 Replication and transcription on a collision course: eukaryotic regulation mechanisms and implications for DNA stability. Frontiers in Genetics 6.

Branzei, D., and I. Psakhye, 2016 DNA damage tolerance. Curr Opin Cell Biol 40: 137–144.

Branzei, D., and B. Szakal, 2016 DNA damage tolerance by recombination: Molecular pathways and DNA structures. DNA Repair (Amst) 44: 68–75.

Branzei, D., F. Vanoli and M. Foiani, 2008 SUMOylation regulates Rad18-mediated template switch. Nature 456: 915–920.

Campos-Doerfler, L., S. Syed and K. H. Schmidt, 2018 Sgs1 Binding to Rad51 Stimulates Homology-Directed DNA Repair in Saccharomyces cerevisiae. Genetics 208: 125–138.

Chang, M., M. Bellaoui, C. Boone and G. W. Brown, 2002 A genome-wide screen for methyl methanesulfonate-sensitive mutants reveals genes required for S phase progression in the presence of DNA damage. Proc Natl Acad Sci U S A 99: 16934–16939.

Chatterjee, N., and G. C. Walker, 2017 Mechanisms of DNA damage, repair, and mutagenesis. Environ Mol Mutagen 58: 235–263.

Chen, J., B. Derfler, A. Maskati and L. Samson, 1989 Cloning a eukaryotic DNA glycosylase repair gene by the suppression of a DNA repair defect in Escherichia coli. Proc Natl Acad Sci U S A 86: 7961–7965.

Collins, S. R., K. M. Miller, N. L. Maas, A. Roguev, J. Fillingham et al., 2007 Functional dissection of protein complexes involved in yeast chromosome biology using a genetic interaction map. Nature 446: 806–810.

Conde, F., D. Ontoso, I. Acosta, A. Gallego-Sanchez, A. Bueno et al., 2010 Regulation of tolerance to DNA alkylating damage by Dot1 and Rad53 in Saccharomyces cerevisiae. DNA Repair (Amst) 9: 1038–1049.

Conde, F., and P. A. San-Segundo, 2008 Role of Dot1 in the response to alkylating DNA damage in Saccharomyces cerevisiae: regulation of DNA damage tolerance by the error-prone polymerases Polzeta/Rev1. Genetics 179: 1197–1210.

Costanzo, M., B. Vandersluis, E. N. Koch, A. Baryshnikova, C. Pons et al., 2016 A global genetic interaction network maps a wiring diagram of cellular function. Science 353.

Daigaku, Y., A. A. Davies and H. D. Ulrich, 2010 Ubiquitin-dependent DNA damage bypass is separable from genome replication. Nature 465: 951–955.

Diebold, M. L., M. Koch, E. Loeliger, V. Cura, F. Winston et al., 2010 The structure of an Iws1/Spt6 complex reveals an interaction domain conserved in TFIIS, Elongin A and Med26. EMBO J 29: 3979–3991.

Downs, J. A., N. F. Lowndes and S. P. Jackson, 2000 A role for Saccharomyces cerevisiae histone H2A in DNA repair. Nature 408: 1001–1004.

Dungrawala, H., K. L. Rose, K. P. Bhat, K. N. Mohni, G. G. Glick et al., 2015 The Replication Checkpoint Prevents Two Types of Fork Collapse without Regulating Replisome Stability. Mol Cell 59: 998–1010.

Fischbeck, J. A., S. M. Kraemer and L. A. Stargell, 2002 SPN1, a conserved gene identified by suppression of a postrecruitment-defective yeast TATA-binding protein mutant. Genetics 162: 1605–1616.

Foster, E. R., and J. A. Downs, 2005 Histone H2A phosphorylation in DNA double-strand break repair. FEBS J 272: 3231–3240.

Gangavarapu, V., S. Prakash and L. Prakash, 2007 Requirement of RAD52 group genes for postreplication repair of UV-damaged DNA in Saccharomyces cerevisiae. Mol Cell Biol 27: 7758–7764.

Gerard, A., E. Segeral, M. Naughtin, A. Abdouni, B. Charmeteau et al., 2015 The integrase cofactor LEDGF/p75 associates with Iws1 and Spt6 for postintegration silencing of HIV-1 gene expression in latently infected cells. Cell Host Microbe 17: 107–117.

Gonzalez-Huici, V., B. Szakal, M. Urulangodi, I. Psakhye, F. Castellucci et al., 2014 DNA bending facilitates the error-free DNA damage tolerance pathway and upholds genome integrity. Embo Journal 33: 327–340.

Guzder, S. N., Y. Habraken, P. Sung, L. Prakash and S. Prakash, 1996a RAD26, the yeast homolog of human Cockayne’s syndrome group B gene, encodes a DNA-dependent ATPase. J Biol Chem 271: 18314–18317.

Guzder, S. N., P. Sung, L. Prakash and S. Prakash, 1996b Nucleotide excision repair in yeast is mediated by sequential assembly of repair factors and not by a pre-assembled repairosome. J Biol Chem 271: 8903–8910.

Hall, B. M., C. X. Ma, P. Liang and K. K. Singh, 2009 Fluctuation analysis CalculatOR: a web tool for the determination of mutation rate using Luria-Delbruck fluctuation analysis. Bioinformatics 25: 1564–1565.

Hanamshet, K., O. M. Mazina and A. V. Mazin, 2016 Reappearance from Obscurity: Mammalian Rad52 in Homologous Recombination. Genes (Basel) 7.

Huang, D., B. D. Piening and A. G. Paulovich, 2013 The preference for error-free or error-prone postreplication repair in Saccharomyces cerevisiae exposed to low-dose methyl methanesulfonate is cell cycle dependent. Mol Cell Biol 33: 1515–1527.

Hung, S. H., R. P. Wong, H. D. Ulrich and C. F. Kao, 2017 Monoubiquitylation of histone H2B contributes to the bypass of DNA damage during and after DNA replication. Proc Natl Acad Sci U S A 114: E2205–E2214.

Hustedt, N., S. M. Gasser and K. Shimada, 2013 Replication Checkpoint: Tuning and Coordination of Replication Forks in S Phase. Genes 4: 388–434.

Kawasaki, Y., and A. Sugino, 2001 Yeast replicative DNA polymerases and their role at the replication fork. Molecules and Cells 12: 277–285.

Kim, J. A., and J. E. Haber, 2009 Chromatin assembly factors Asf1 and CAF-1 have overlapping roles in deactivating the DNA damage checkpoint when DNA repair is complete. Proc Natl Acad Sci U S A 106: 1151–1156.

Koc, A., L. J. Wheeler, C. K. Mathews and G. F. Merrill, 2004 Hydroxyurea arrests DNA replication by a mechanism that preserves basal dNTP pools. J Biol Chem 279: 223–230.

Krogan, N. J. M., M. Kim, S. H. Ahn, G. Zhong, M. S. Kobor et al., 2002 RNA polymerase II elongation factors of Saccharomyces cerevisiae: a targeted proteomics approach. Mol Cell Biol 22: 6979–6992.

Kubota, T., D. A. Stead, S. Hiraga, S. Ten Have and A. D. Donaldson, 2012 Quantitative proteomic analysis of yeast DNA replication proteins. Methods 57: 196–202.

Lea, D. E., and C. A. Coulson, 1949 The Distribution of the Numbers of Mutants in Bacterial Populations. Journal of Genetics 49: 264–285.

Lee, K. Y., and K. Myung, 2008 PCNA modifications for regulation of post-replication repair pathways. Mol Cells 26: 5–11.

Li, S., A. R. Almeida, C. A. Radebaugh, L. Zhang, X. Chen et al., 2018 The elongation factor Spn1 is a multi-functional chromatin binding protein. Nucleic Acids Res 46: 2321–2334.

Liu, Z., Z. Zhou, G. Chen and S. Bao, 2007 A putative transcriptional elongation factor hIws1 is essential for mammalian cell proliferation. Biochem Biophys Res Commun 353: 47–53.

Longo, V. D., and P. Fabrizio, 2012 Chronological Aging in Saccharomyces cerevisiae. Aging Research in Yeast 57: 101–121.

Madia, F., C. Gattazzo, P. Fabrizio and V. D. Longo, 2007 A simple model system for age-dependent DNA damage and cancer. Mech Ageing Dev 128: 45–49.

Madia, F., M. Wei, V. Yuan, J. Hu, C. Gattazzo et al., 2009 Oncogene homologue Sch9 promotes age-dependent mutations by a superoxide and Rev1/Polzeta-dependent mechanism. J Cell Biol 186: 509–523.

Mcdonald, S. M., D. Close, H. Xin, T. Formosa and C. P. Hill, 2010 Structure and biological importance of the Spn1-Spt6 interaction, and its regulatory role in nucleosome binding. Mol Cell 40: 725–735.

Meas, R., M. J. Smerdon and J. J. Wyrick, 2015 The amino-terminal tails of histones H2A and H3 coordinate efficient base excision repair, DNA damage signaling and postreplication repair in Saccharomyces cerevisiae. Nucleic Acids Res 43: 4990–5001.

Memisoglu, A., and L. Samson, 2000 Base excision repair in yeast and mammals. Mutat Res 451: 39–51.

Mitchel, K., K. Lehner and S. Jinks-Robertson, 2013 Heteroduplex DNA position defines the roles of the Sgs1, Srs2, and Mph1 helicases in promoting distinct recombination outcomes. PLoS Genet 9: e1003340.

Mullen, J. R., F. S. Nallaseth, Y. Q. Lan, C. E. Slagle and S. J. Brill, 2005 Yeast Rmi1/Nce4 controls genome stability as a subunit of the Sgs1-Top3 complex. Mol Cell Biol 25: 4476–4487.

Odell, I. D., S. S. Wallace and D. S. Pederson, 2013 Rules of engagement for base excision repair in chromatin. J Cell Physiol 228: 258–266.

Parrella, E., and V. D. Longo, 2008 The chronological life span of Saccharomyces cerevisiae to study mitochondrial dysfunction and disease. Methods 46: 256–262.

Pfister, S. X., S. Ahrabi, L. P. Zalmas, S. Sarkar, F. Aymard et al., 2014 SETD2-dependent histone H3K36 trimethylation is required for homologous recombination repair and genome stability. Cell Rep 7: 2006–2018.

Polo, S. E., and G. Almouzni, 2015 Chromatin dynamics after DNA damage: The legacy of the access-repair-restore model. DNA Repair (Amst) 36: 114–121.

Prakash, L., and S. Prakash, 1977 Isolation and characterization of MMS-sensitive mutants of Saccharomyces cerevisiae. Genetics 86: 33–55.

Prakash, S., R. E. Johnson and L. Prakash, 2005 Eukaryotic translesion synthesis DNA polymerases: specificity of structure and function. Annu Rev Biochem 74: 317–353.

Prakash, S., and L. Prakash, 2000 Nucleotide excision repair in yeast. Mutat Res 451: 13–24.

Pujari, V., C. A. Radebaugh, J. V. Chodaparambil, U. M. Muthurajan, A. R. Almeida et al., 2010 The transcription factor Spn1 regulates gene expression via a highly conserved novel structural motif. J Mol Biol 404: 1–15.

Skoneczna, A., A. Kaniak and M. Skoneczny, 2015 Genetic instability in budding and fission yeast-sources and mechanisms. Fems Microbiology Reviews 39: 917–967.

Stone, J. E., S. A. Lujan, T. A. Kunkel and T. A. Kunkel, 2012 DNA polymerase zeta generates clustered mutations during bypass of endogenous DNA lesions in Saccharomyces cerevisiae. Environ Mol Mutagen 53: 777–786.

Symington, L. S., R. Rothstein and M. Lisby, 2014 Mechanisms and regulation of mitotic recombination in Saccharomyces cerevisiae. Genetics 198: 795–835.

Torres-Ramos, C. A., R. E. Johnson, L. Prakash and S. Prakash, 2000 Evidence for the involvement of nucleotide excision repair in the removal of abasic sites in yeast. Mol Cell Biol 20: 3522–3528.

Ulrich, H. D., 2011 Timing and spacing of ubiquitin-dependent DNA damage bypass. FEBS Lett 585: 2861–2867.

Van Attikum, H., O. Fritsch and S. M. Gasser, 2007 Distinct roles for SWR1 and INO80 chromatin remodeling complexes at chromosomal double-strand breaks. EMBO J 26: 4113–4125.

Vijg, J., and Y. Suh, 2013 Genome instability and aging. Annu Rev Physiol 75: 645–668.

Wyatt, M. D., and D. L. Pittman, 2006 Methylating agents and DNA repair responses: Methylated bases and sources of strand breaks. Chem Res Toxicol 19: 1580–1594.

Xu, X., S. Blackwell, A. Lin, F. Li, Z. Qin et al., 2015 Error-free DNA-damage tolerance in Saccharomyces cerevisiae. Mutat Res Rev Mutat Res 764: 43–50.

Yang, Y., D. A. Gordenin and M. A. Resnick, 2010 A single-strand specific lesion drives MMS-induced hyper-mutability at a double-strand break in yeast. DNA Repair (Amst) 9: 914–921.

Yoh, S. M., H. Cho, L. Pickle, R. M. Evans and K. A. Jones, 2007 The Spt6 SH2 domain binds Ser2-P RNAPII to direct Iws1-dependent mRNA splicing and export. Genes Dev 21: 160–174.

Yoh, S. M., J. S. Lucas and K. A. Jones, 2008 The Iws1:Spt6:CTD complex controls cotranscriptional mRNA biosynthesis and HYPB/Setd2-mediated histone H3K36 methylation. Genes Dev 22: 3422–3434.

Zeman, M. K., and K. A. Cimprich, 2014 Causes and consequences of replication stress. Nat Cell Biol 16: 2–9.

Zhang, L., A. G. Fletcher, V. Cheung, F. Winston and L. A. Stargell, 2008 Spn1 regulates the recruitment of Spt6 and the Swi/Snf complex during transcriptional activation by RNA polymerase II. Mol Cell Biol 28: 1393–1403.

